# Dopaminergic genes are associated with both directed and random exploration

**DOI:** 10.1101/357251

**Authors:** Samuel J. Gershman, Bastian Greshake Tzovaras

**Affiliations:** Department of Psychology and Center for Brain Science, Harvard University; Lawrence Berkeley National Laboratory, Berkeley, CA, USA

**Author notes:** Address for correspondence: Samuel Gershman Department of Psychology Harvard University 52 Oxford St., room 295.05 Cambridge, MA 02138.

**Keywords:** explore-exploit dilemma, reinforcement learning, Bayesian inference

## Abstract

In order to maximize long-term rewards, agents must balance exploitation (choosing the option with the highest payoff) and exploration (gathering information about options that might have higher payoffs). Although the optimal solution to this trade-off is intractable, humans make use of two effective strategies: selectively exploring options with high uncertainty (directed exploration), and increasing the randomness of their choices when they are more uncertain (random exploration). Using a task that independently manipulates these two forms of exploration, we show that single nucleotide polymorphisms related to dopamine are associated with individual differences in exploration strategies. Variation in a gene linked to prefrontal dopamine (COMT) predicted the degree of directed exploration, as well as the overall randomness of responding. Variation in a gene linked to striatal dopamine (DARPP-32) predicted the degree of both directed and random exploration. These findings suggest that dopamine makes multiple contributions to exploration, depending on its afferent target.

## Introduction

Any agent seeking to maximize long-term rewards faces an exploration-exploitation dilemma: should it exploit the option with the highest observed payoff, or should it explore other options that might yield higher payoffs? This problem is, in general, intractable, but computer scientists have developed simple and provably effective strategies (Ghavamzadeh et al., 2015). Most of these strategies fall into one of two classes. The first class consists of directed strategies such as Upper Confidence Bound (UCB; Auer et al., 2002), which add an “uncertainty bonus” to the value estimate of each action, selecting action *a*_*t*_ on trial t according to:

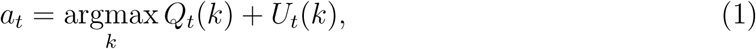

where *U*_*t*_(*k*) is the uncertainty bonus. In one version of this model previously used to study humans (Gershman, 2018a), the uncertainty bonus is proportional to the posterior standard deviation *σ*_*t*_(*k*) (Srinivas et al., 2010), where the posterior over values is updated using Bayes’ rule (see Materials and Methods). The second class consists of random strategies, such as Thompson Sampling (Thompson, 1933), which increase randomness when value estimates are uncertain. Specifically, Thompson sampling draws random values from the posterior and then chooses greedily with respect to these random values. Both directed and random strategies formalize the idea that an agent should gather information about actions when it is uncertain about them, in order to more quickly discover which actions are the most rewarding.

Humans appear to use both directed and random exploration strategies (Gershman, 2018a; Krueger et al., 2017; Wilson et al., 2014). These strategies appear to rely on different neural systems (Warren et al., 2017; Zajkowski et al., 2017), follow different developmental trajectories (Schulz et al., 2018; Somerville et al., 2017), and can be independently manipulated (Gershman, 2018b). An important gap in this emerging picture concerns the role of the neuromodulator dopamine, which plays a central role in reinforcement learning theories (Glimcher, 2011). Some theories hypothesize that dopamine responses to novelty reflect an uncertainty bonus consistent with directed exploration strategies (Kakade and Dayan, 2002), whereas other theories hypothesize that dopamine controls the degree of random exploration through gain modulation of striatal neurons (Friston et al., 2012; Humphries et al., 2012). These two roles are not intrinsically in conflict, but no research has yet tried to disentangle them empirically.

We pursue this question by studying single nucleotide genetic polymorphisms related to dopamine function. A polymorphism in the gene encoding catechol-O-methyltransferase (COMT), an enzyme that breaks down dopamine, has been shown to have a relatively selective effect on dopamine transmission in prefrontal cortex (Slifstein et al., 2008). A previous study by Frank et al. (2009) found that individuals with the methionine-encoding allele, putatively associated with higher prefrontal dopamine levels, exhibited greater levels of directed exploration. However, this study did not try to disentangle directed and random exploration strategies, and its experimental design was not ideal for this purpose.

In this paper, we use a two-armed bandit task specifically designed to disentangle directed and random exploration strategies (Gershman, 2018b). Subjects receive information about the riskiness of each arm; a “risky” arm pays off stochastically, whereas a “safe” arm pays off deterministically (Figure 1). By allowing each arm to be either risky or safe in a block of trials, the design allows us to separate the effects of directed and random exploration (Figure 2). Because directed exploration is based on the addition of uncertainty bonuses, the critical variable is the *relative* uncertainty between the two arms. Thus, we can isolate the effect of directed exploration by comparing trials in which one option is risky and the other is safe. The uncertainty bonus accruing to the risky option will cause choice probability, plotted as a function of the difference in value estimates between the two arms, to shift its intercept (i.e., the indifference point). To illustrate, if option 1 has a larger uncertainty than option 2, then the choice probability function will shift to the left, reflecting an increased preference for option 1.

**Figure 1:**
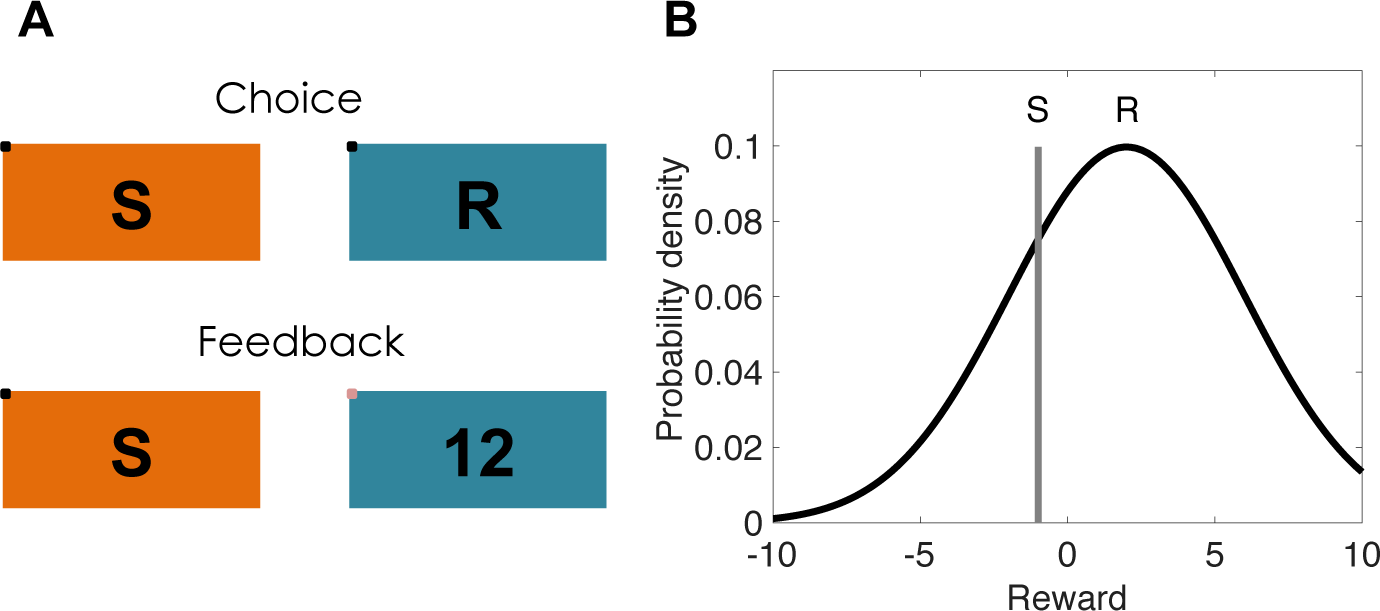
Task design. (Left) On each trial, subjects choose between two options and receive reward feedback in the form of points. Safe options are denoted by “S” and risky options are denoted by “R.” On each block, one or both of the options may be safe or risky. (Right) The rewards for risky options are drawn from a Gaussian distribution that remains constant during each block. The rewards for safe options are deterministic. Both the mean for the risky option and the reward value of the safe option are drawn from a zero-mean Gaussian distribution that is resampled at each block transition.

**Figure 2:**
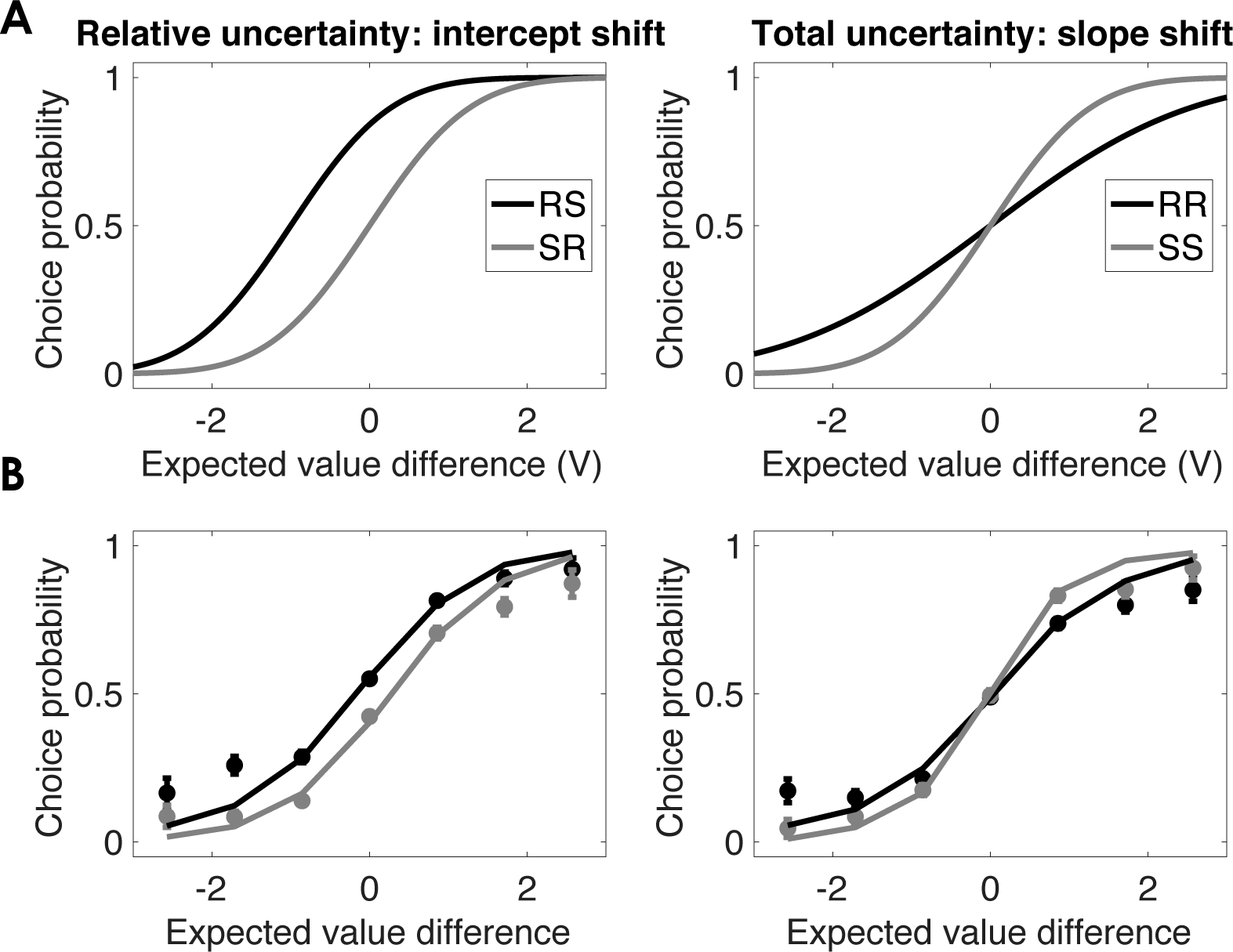
Choice probability functions. (A) Illustration of how the probability of choosing option 1 changes as a function of the experimental condition and form of uncertainty. V represents the difference between the expected value of option 1 and the expected value of option 2. (B) Empirical choice probability functions. Lines show probit model fits. Error bars denote within-subject standard error of the mean.

In contrast, random exploration is based on increasing stochasticity in proportion to *total* uncertainty across the arms. This will manifest as a slope change in the choice probability function, without altering the intercept. To illustrate, if both options are safe, the choice probability function will become steeper compared to if both options are risky, such that the change in choice probability as a function of a unit change in value difference is greater.

We leverage this task to re-examine the role of COMT in different exploration strategies. Although the study of Frank and colleagues (Frank et al., 2009) suggested that COMT mediates directed exploration, another prominent theory of COMT suggests that it controls the gain of prefrontal neurons (Durstewitz and Seamans, 2008), which would lend naturally lend itself to the modulation of random exploration. Thus, prior work indicates the plausibility of COMT involvement in both directed and random exploration.

In addition to COMT, our study also examines a polymorphism in the gene encoding the DARPP-32 phosphoprotein, which controls the excitability and plasticity of neurons in the striatum that receive dopamine (Fienberg et al., 1998; Schiffmann et al., 1998). Individuals with two copies of the T allele have greater DARPP-32 mRNA expression in striatum, which has been linked to differences in learning from rewards, but not to differences in exploration strategy (Frank et al., 2009). Other models have implicated striatal dopamine modulation in the decision process. Based on biophysical modeling of striatal neurons (Humphries et al., 2012), we hypothesized that DARPP-32 would play a role in random exploration. Specifically, increased levels of DARPP-32 in the striatum should increase the gain of neurons expressing the D1 receptor, which would cause increased selection of high-value options (i.e., reduced random exploration). On the other hand, pharmacological (Stopper et al., 2013) and electrophysiological (Naudé et al., 2018) experiments have suggested that striatal dopamine transmission mediates a bias towards risky choice, consistent with an effect on directed exploration. Thus, DARPP-32, like COMT, could plausibly be involved in both directed and random exploration.

## Materials and Methods

### Participants

We recruited participants from openSNP, an open database where individuals publish raw data of their existing direct-to-consumer (DTC) genetic tests as performed by 23andMe, FamilyTreeDNA or AncestryDNA (Greshake et al., 2014). We emailed 867 members of openSNP who had published genetic data that include the variants rs4680 in the COMT gene and rs907094 in the DARPP-32 gene. Of those contacted, 121 members participated in the experimental task through an online interface. The experiment was approved by the Harvard University Institutional Review Board.

We excluded 5 participants who had very low accuracy (selecting the optimal choice less than 55% of the time). The remaining 116 participants (60 females) had an age range of 15-90 (median: 41) and were 98% Caucasian.

### Genotyping

The genetic tests that analyzed these variants were performed in the laboratories of 23andMe and/or AncestryDNA using the companies’ respective DNA micro arrays. Because the genotyping methods of those companies are not publicly available, we cannot report them here. Neither genotype deviated significantly from Hardy-Weinberg equilibrium, according to a chi-squared test.

In order to simplify the statistical analysis and ensure comparability with previous results, we clustered genotypes into two groups using the same clusters as Frank and colleagues (Frank et al., 2009), which in this case also maps onto putative differences in regionally specific dopamine transmission. The COMT genotype was clustered into Val/Val (putatively low prefrontal dopamine transmission, *N* = 27) and Val/Met + Met/Met (putatively high prefrontal dopamine transmission, *N* = 89) groups. The DARPP-32 genotypes was clustered into C/C + C/T (putatively low striatal dopamine transmission, *N* = 48) and T/T (putatively high striatal dopamine transmission, *N* = 68) groups. No age differences were found between the genotype clusters for either polymorphism (two-sample t-test: *p* = 0.11 for COMT, *p* = 0.66 for DARPP-32). Likewise, no gender differences were found between the genotype clusters for either polymorphism (binomial test: *p* = 0.5 for COMT, *p* = 0.52 for DARPP-32). The regression results reported below were qualitatively unchanged when age and gender were included as covariates.

### Experimental Procedure

Participants played 30 two-armed bandits, each for one block of 10 trials. On each trial, participants chose one of the arms and received reward feedback (points). They were instructed to choose the “slot machine” (corresponding to an arm) that maximizes their total points. On each block, the mean reward *μ*(*k*) for each arm was drawn from a Gaussian with mean 0 and variance 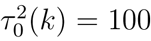. The arms were randomly designated “safe” or “risky,” indicated by an S or R (respectively), and these designations were randomly resampled after a block transition. When participants chose the risky arm, they received stochastic rewards drawn from a Gaussian with mean *μ*(*R*) and variance *τ*^2^(*R*) = 16. When participants chose the safe arm, they received a reward of *μ*(*S*). We use compound notation to denote trial type: for example, “SR” denotes a trial in which arm 1 is safe and arm 2 is risky.

The exact instructions for participants were as follows:

> *In this task, you have a choice between two slot machines, represented by colored buttons. When you click one of the buttons, you will win or lose points. Choosing the same slot machine will not always give you the same points, but one slot machine is always better than the other. Your goal is to choose the slot machine that will give you the most points. After making your choice, you will receive feedback about the outcome. Sometimes the machines are “safe” (always delivering the same feedback), and sometimes the machines are “risky” (delivering variable feedback). Before you make a choice, you will get information about each machine: “S” indicates SAFE, “R” indicates RISKY. Note that safe/risky is independent of how rewarding a machine is: a risky machine may deliver more points on average than a safe machine, and vice versa. You cannot predict how good a machine is based on whether it is safe or risky. You will play 30 games, each with a different pair of slot machines. Each game will consist of 10 trials*.

### Modeling

To derive estimates of expected value and uncertainty, we assume that subjects approximate an ideal Bayesian learner. Given the Gaussian distributional structure underlying our task, the posterior over the value of arm *k* is Gaussian with mean *Q*_*t*_(*k*) and variance 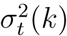:

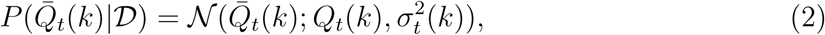

where *Q̄*_*t*_(*k*) is the true value of arm *k* and 𝒟 denotes the history of choices and rewards. The sufficient statistics can be recursively updated using the Kalman filtering equations:

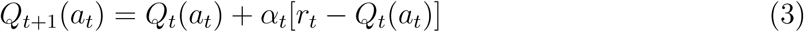

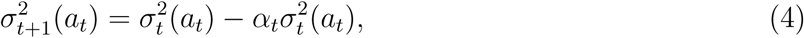

where *a*_*t*_ is the chosen arm, *r*_*t*_ is the received reward, and the learning rate *α*_*t*_ is given by:

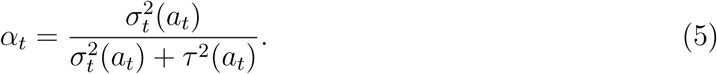

Note that only the chosen option’s mean and variance are updated after each trial.^1^

The initial values were set to the prior means, *Q*_1_(*k*) = 0 for all *k*, and the initial variances were set to the prior variance, 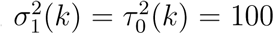. The value of *τ*^2^ was set to 16 (its true value) for risky options, and to 0.00001 for safe options (to avoid numerical issues, the variance was not set exactly to 0). Although the Kalman filter is an idealization of human learning, it has been shown to account well for human behavior in bandit tasks (Daw et al., 2006b; Gershman, 2018a; Schulz et al., 2015; Speekenbrink and Konstantinidis, 2015).

Previous work (Gershman, 2018a) showed that Thompson sampling can be formalized as a probit regression model of choice:

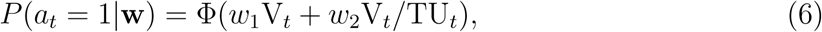

where *V*_*t*_ = *Q*_*t*_(1) – *Q*_*t*_(2) is the estimated value difference, 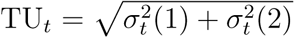 denotes total uncertainty, and Φ(·) is the cumulative distribution function of the standard Gaussian distribution (mean 0 and variance 1). In contrast, UCB, assuming Gaussian noise corrupting the value estimates, can be formalized as the following probit regression model:

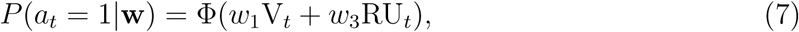

where RU_*t*_ = *σ*_*t*_(1) – *σ*_*t*_(2) denotes relative uncertainty. We modeled a hybrid of the two strategies by including both random and directed terms:

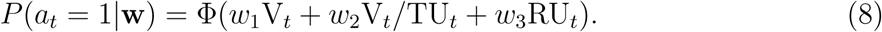

We also analyzed choices as a function of experimental condition (i.e., RS, SR, RR, SS):

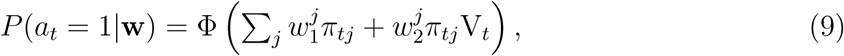

where and *σ*_*tj*_ = 1 if trial *t* is assigned to condition *j*, and 0 otherwise. We refer to the *w*_1_ terms as intercepts and the *w*_2_ terms as slopes. For the full model, we specify genotype clusters as fixed effects that interact with condition type, such that was have separate intercepts and slopes for each genotype cluster. These parameters were estimated using a generalized linear mixed-effects model implemented in Matlab (version R2017b).

## Results

We analyzed the choice behavior of 116 individuals participating in the risky-safe two-armed bandit described above (see Materials and Methods for more details). Subjects were fairly accurate on the task, choosing the optimal action on average 73% of the time (±0.6 standard error of the mean). Using mixed-effects probit regression, we first sought to replicate the results of a previous study using this task (Gershman, 2018b). Consistent with that study, we found a significant effect of the relative uncertainty manipulation on the intercept parameter [RS vs. SR, *F*(1, 34752) = 40.46, *p* < 0.001], but not on the slope parameter (*p* = 0.08), and a significant effect of the total uncertainty manipulation on the slope parameter [SS vs. RR, *F*(1, 34752) = 71.65, *p* < 0.001], but not on the intercept parameter (*p* = 0.25). Thus, our results confirm the theoretical predictions of a hybrid model (Gershman, 2018b) in which manipulating relative uncertainty affects directed exploration, while manipulating total uncertainty affects random exploration.

To verify that both random and directed exploration strategies are needed to explain the behavior, we carried out model comparison using standard model comparison metrics (the Bayesian information criterion and the Akaike information criterion).^2^ The models varied in which components they included in the probit regression: value (*V*), a random exploration component (*V*/*TU*), a directed exploration component (*RU*), or both random and directed components. The technical definitions of these components can be found in the Materials and Methods. Among these models, the full model with both exploration components was strongly favored (Table 1). In this model, both the random and directed regression weights were significantly greater than 0 (both *p* < 0.00001). The quantitative support for a random exploration component is particularly important, because differences in slope could conceivably arise from poorer learning in the RR condition compared to the SS condition.

**Table 1:** Model comparison results. Each row corresponds to a probit regression model, with the first column indicating the terms included in the model (see Materials and Methods for a technical description of these models). BIC = Bayesian information criterion; AIC = Akaike information criterion. Lower values indicate stronger model evidence.

| Model | BIC | AIC | pseudo-R<sup>2</sup> |
| --- | --- | --- | --- |
| V | 13493 | 13478 | 0.245 |
| V/TU | 17262 | 17247 | 0.075 |
| V + RU | 12874 | 12836 | 0.317 |
| V + RU + V/TU | 12689 | 12621 | 0.332 |

We next asked how genetic variation in dopamine neuromodulation affected the different exploration strategies, using the genotype clusters from a previous study (Frank et al., 2009). As shown in Figure 3, variation in COMT had two distinct effects. First, subjects with the Met allele (associated with higher levels of prefrontal dopamine) showed a larger effect of the relative uncertainty manipulation on the intercept parameter [*F*(1, 34744) = 8.06, *p* < 0.005]. In other words, subjects with putatively higher levels of prefrontal dopamine exhibited a directed exploration strategy that was more sensitive to the relative uncertainty manipulation, a pattern that can be seen visually in Figure 4 (top panels). We did not find a significant interaction between COMT and total uncertainty for the slope parameter (*p* = 0.37), indicating that COMT did not influence the degree of random exploration.

**Figure 3:**
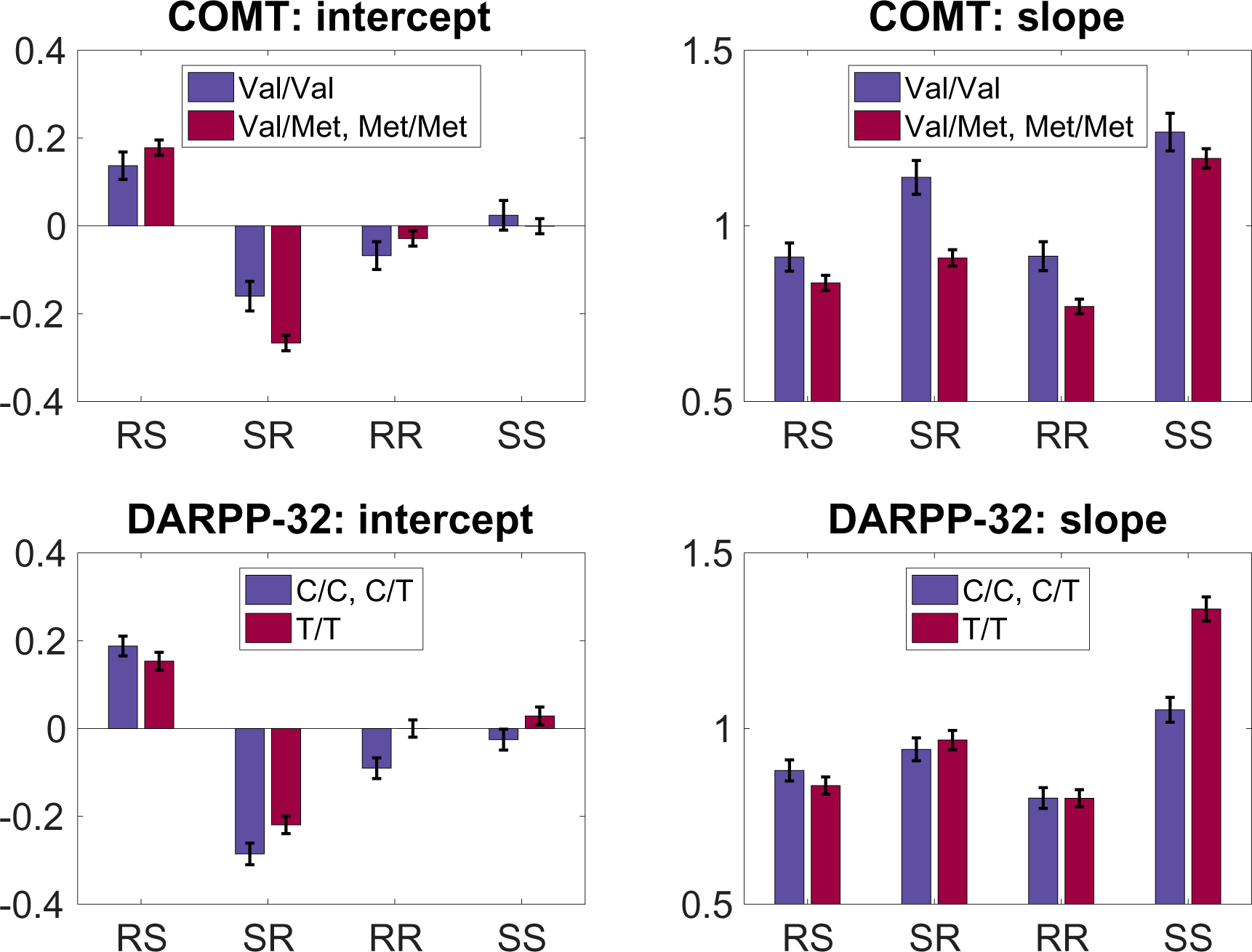
Regression results. Slope and intercept parameter estimates broken down by genotype cluster and experimental condition. Error bars represent standard errors.

**Figure 4:**
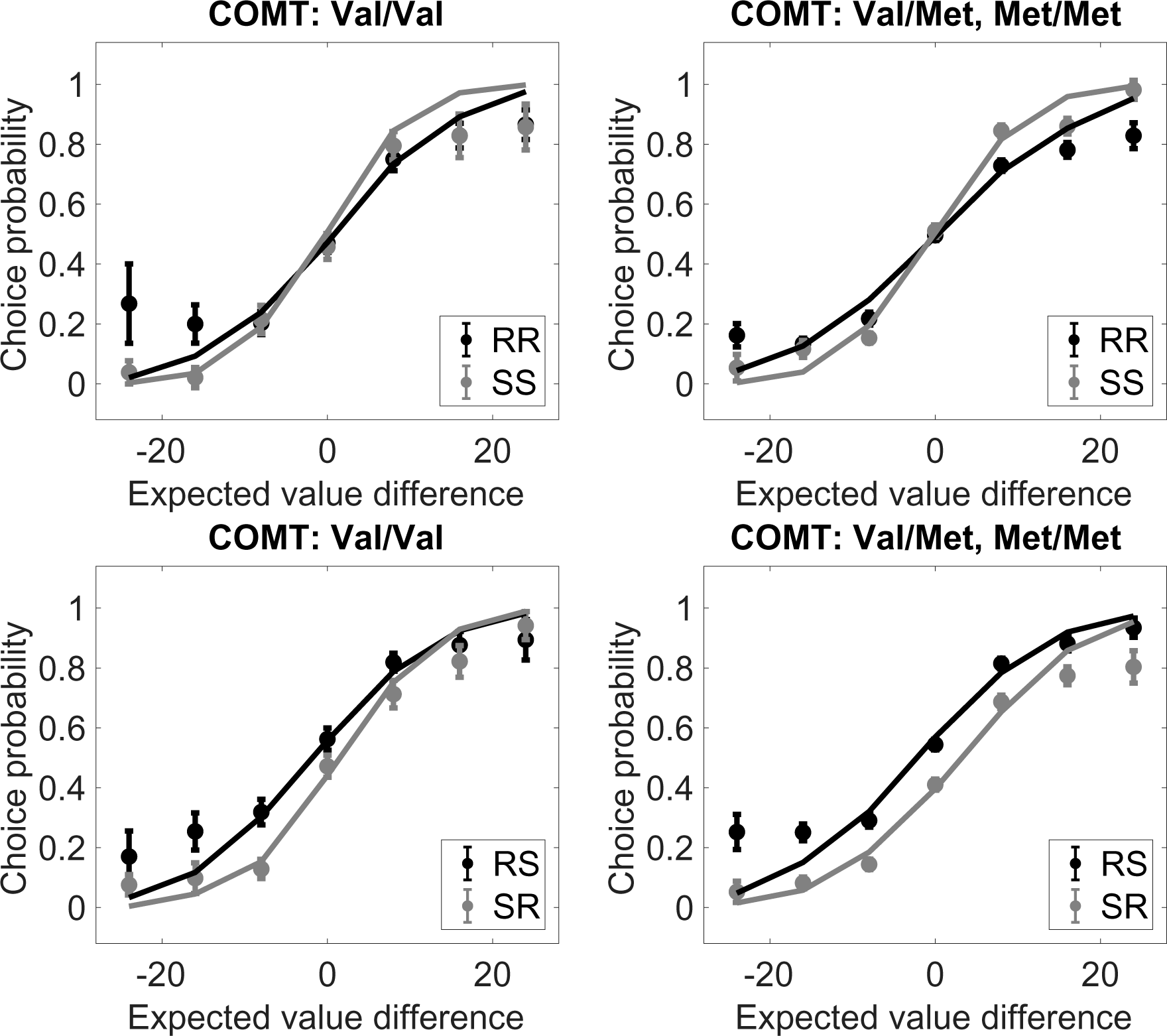
Choice probability functions: COMT. Choice probabilities broken down by genotype cluster and experimental condition. Lines show probit model fits. Error bars represent within-subject standard error of the mean.

In addition to its effect on directed exploration, variation in COMT was also associated with an overall change in slope across experimental conditions. Specifically, carriers of the Met allele had a lower slope compared to Val/Val carriers, indicating that choice was more random (independent of relative or total uncertainty) for subjects with putatively higher levels of prefrontal dopamine. Note that we reserve the term “random exploration” to refer to the effect of total uncertainty on choice randomness, which is distinct from the overall randomness across conditions, akin to what is captured by the inverse temperature in a standard softmax policy. Thus, Met carriers were overall more random without modulating their randomness as a function of total uncertainty.

Variation in DARPP-32 also had two distinct effects (Figure 3). First, there was a significant interaction between genotype cluster and the total uncertainty manipulation for the slope parameter [*F*(1, 34744) = 21.33, *p* < 0.0001], such that T/T carriers (associated with higher striatal D1 transmission) had a higher difference in slope (less randomness) between the SS and RR conditions, a finding confirmed visually in the choice probability functions (Figure 5). As shown in Figure 3, this was primarily driven by higher slope in the SS condition for T/T carriers.

**Figure 5:**
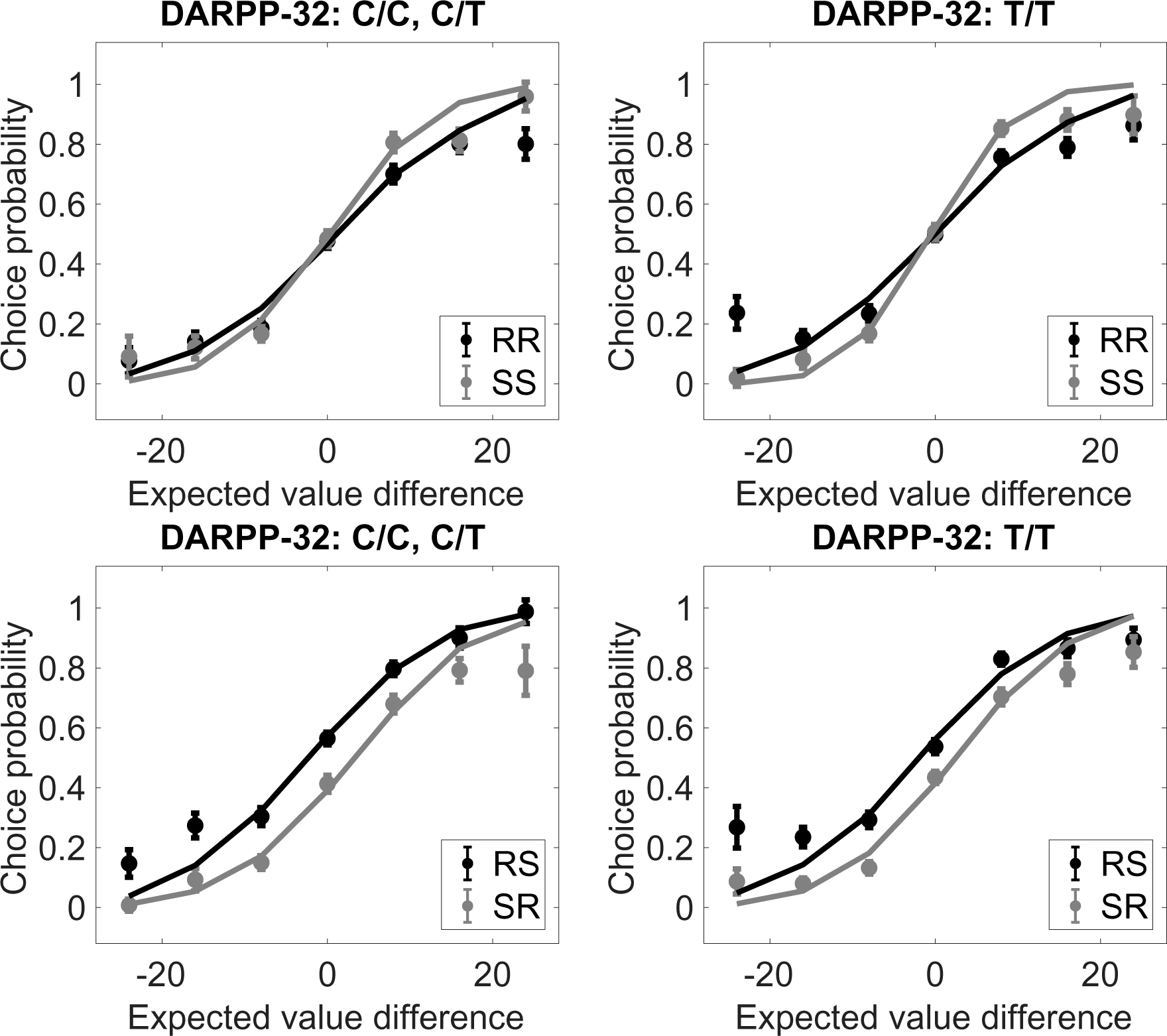
hoice probability functions: DARPP-32. Choice probabilities broken down by genotype cluster and experimental condition. Lines show probit model fits. Error bars represent within-subject standard error of the mean.

Second, there was a significant interaction between genotype cluster and the relative uncertainty manipulation for the intercept parameter [*F*(1, 34744) = 5.24, *p* < 0.05], such that T/T carriers showed a reduced effect of relative uncertainty on directed exploration. Thus, whereas putatively higher prefrontal dopamine predicted a larger “uncertainty bonus,” this bonus was smaller for putatively higher striatal dopamine.

In addition to our condition-based analysis, we performed a model-based analysis of the interaction between genotype cluster and the three computational regressors (V, RU, V/TU). As shown in Figure 6, these results largely recapitulate the condition-based analysis. First, inverse temperature (the parameter estimate for V) was significantly higher for COMT Val/Val [*F*(1, 34744) = 4.07, *p* < 0.05]. Second, the effect of relative uncertainty was significantly lower for COMT Val/Val [*F*(1, 34744) = 5.03, *p* < 0.05]. Third, the effect of total uncertainty was significantly higher for DARPP-32 T/T [*F*(1, 34744) = 20.18, *p* < 0.0001]. The one discrepancy with the condition-based analysis was the absence of a significant RU effect for DARPP-32 (*p* = 0.56).

**Figure 6:**
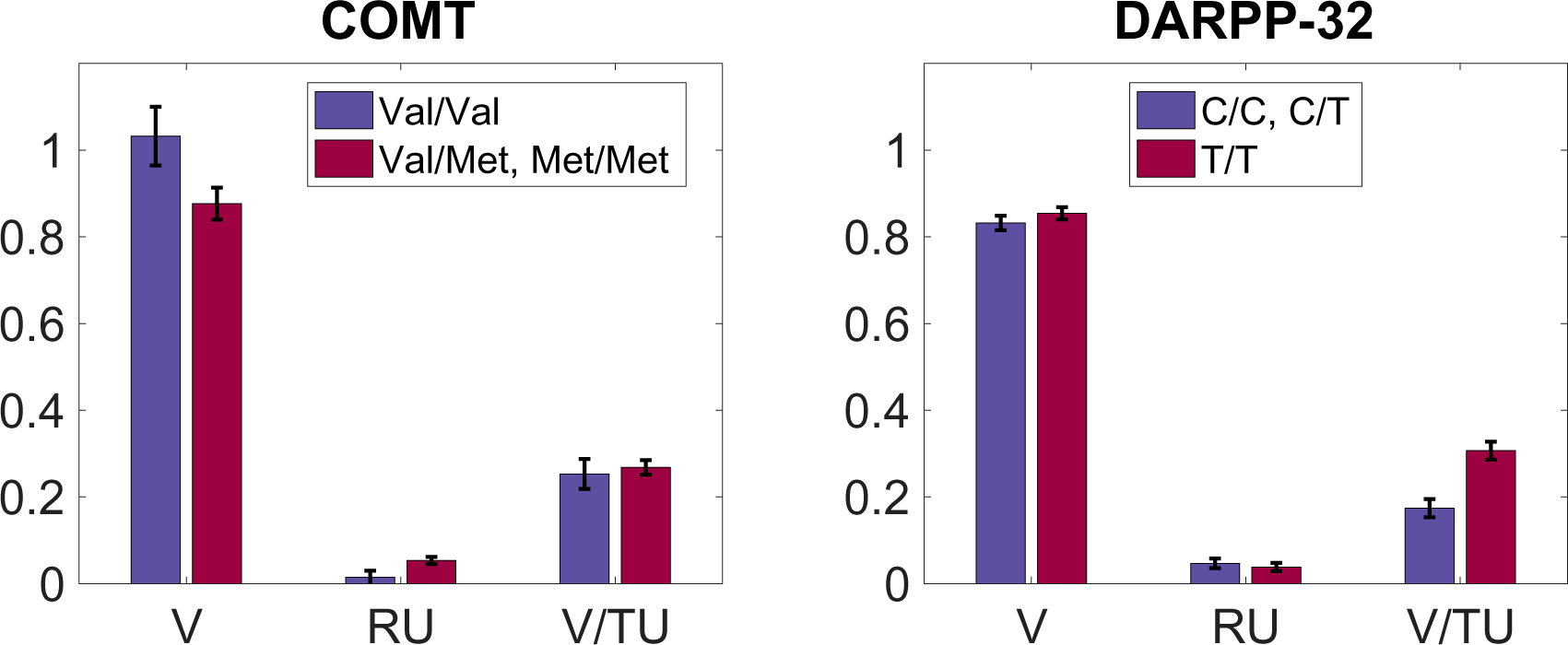
Model-based regression results. Parameter estimates for value (V), relative uncertainty (RU), and value normalized by total uncertainty (V/TU), broken down by genotype cluster and experimental condition. Error bars represent standard errors.

## Discussion

The findings of this study indicate that two dopaminergic genes (COMT and DARPP-32) have multifaceted roles in the orchestration of exploration strategies. A single nucleotide polymorphism in the COMT gene, which has been found to influence prefrontal dopamine transmission, was associated with changes in directed exploration, such that greater putative prefrontal dopamine transmission (carriers of the Met allele) produced a larger effect of relative uncertainty on directed exploration, consistent with the idea that COMT mediates an “uncertainty bonus” for exploration (Frank et al., 2009). In addition, Met carriers appeared to choose more randomly across all conditions (i.e., have a lower inverse temperature), independent of relative and total uncertainty. By contrast, a single nucleotide polymorphism in the DARPP-32 gene, which has been found to influence striatal dopamine transmission, was associated with changes in both directed and random exploration: T/T homozygotes, for whom striatal dopamine transmission is putatively higher, exhibited reduced directed exploration (a smaller uncertainty bonus; though this finding was not recapitulated by the model-based analysis) as well as reduced random exploration (less uncertainty-induced choice stochasticity).

These findings are consistent with several existing theoretical ideas about prefrontal dopamine and COMT. Durstewitz and Seamans (2008) proposed that prefrontal dopamine (regulated by COMT) determines the shape of the energy landscape for network activity. When dopamine levels are low, dopamine primarily binds to D2 receptors, which flattens energy barriers between attractors, allowing the network to be more flexible. When dopamine levels are relatively high, dopamine primarily binds to D1 receptors, which deepens energy barriers, allowing the network to maintain a tonic activity state. When dopamine levels are even higher, the network returns to a D2-dominated state characterized by shallow energy barriers. Because of this non-monotonicity, COMT Met carriers could be more flexible or less flexible than Val carriers (Durstewitz and Seamans, 2008, argue that they will typically be less flexible). If we view choice stochasticity as one form of flexibility, then the effect of COMT on inverse temperature across conditions suggests that the Val/Val homozygotes are in a D1-dominated state (low flexibility) while the Met carriers are in a D2-dominated state (high flexibility). This appears to be at odds with the argument of Durstewitz and Seamans (and experimental data reported by Colzato et al., 2010), although the inverted-U dependence of flexibility on dopamine makes this argument a bit slippery. Bilder et al. (2004), by contrast, have argued that the Met allele is associated with lower flexibility, due to its combined effects on cortical and striatal dopamine activity.

Our finding that COMT Met carriers have a larger uncertainty bonus is related to prior work on dopaminegic control of risk preferences. Onge et al. (2011) found that blockade of prefrontal D1 receptors decreased the preference for a risky option in rats, and microdialysis methods using the same task demonstrated that prefrontal dopamine levels track in the risky reward probabilities (Onge et al., 2012). Risk-seeking can also be increased by chemogenetic activation of the mesocortical pathway (Verharen et al., 2018). These findings converge with evidence that prefrontal dopamine, and the COMT gene in particular, plays an important role in uncertainty-directed exploration (Blanco et al., 2015; Frank et al., 2009).

Turning now to striatal dopamine and DARPP-32, our results also fit with some existing theories. Using detailed biophysical simulations, Humphries et al. (2012) argued that exploration could be controlled by D1 receptor-mediated changes in the excitability of striatal neurons; higher levels of D1 receptor activation result in an increase of the input-output gain, which in turn decreases the entropy (i.e., randomness) of the action selection probability distribution. This view broadly fits with our observation that DARPP-32 T/T homozygotes show less random exploration, although Humphries et al. (2012) did not directly address the question of uncertainty-based exploration. More closely related is the Bayesian framework developed by Friston et al. (2012), according to which dopamine encodes precision (inverse variance), ultimately acting as an uncertainty-based gain modulation mechanism (see also Friston et al., 2015). Our finding that DARPP-32 T/T homozygotes showed a reduced uncertainty bonus is harder to reconcile with some previous pharmacological and electrophysiological findings, which have associated striatal dopamine with a *stronger* preference for risk (Naude et al., 2018; Stopper et al., 2013).

Previous modeling of DARPP-32 in reinforcement learning tasks has suggested that it influences learning from gains (relative to losses) via its control over D1-mediated plasticity of striatal medium spiny neurons (Frank et al., 2009). Here, we have not modeled differential learning from gains and losses; we assumed learning followed a Bayesian ideal observer, and modeled genetic effects on the decision process, motivated by data suggesting that DARPP-32 can control the excitability of striatal neurons (Fienberg et al., 1998; Schiffmann et al., 1998). Disentangling learning and decision processes is intrinsically challenging because different modeling assumptions can mimic each other. In particular, an agent that learns more from positive than from negative prediction errors will become risk seeking, producing behavior resembling directed exploration (Niv et al., 2012). This means that the same behavior could be explained by a differential learning rate or an explicit risk preference. Applied to our data, this would imply that DARPP-32 T/T homozygotes would have a *lower* learning rate for gains than losses, since the relative uncertainty effect is smaller in that genotype cluster compared to C allele carriers. This would appear to be the opposite of the finding from Frank and colleagues. Thus, some discrepancies across studies remain to be resolved.

Our study has a number of limitations. First, the sample size is small relative to standards in behavioral genetics (Hewitt, 2012). Replication with a larger sample size is necessary to ensure that our effects are reliable. Second, while this study suggests distinct computational roles for prefrontal and striatal dopamine in uncertainty-based exploration, it is also important to acknowledge that prefrontal cortex and striatum interact in complex ways, with different dopamine regulatory dynamics (Bilder et al., 2004). An important task for future research is to develop a more mechanistic and biologically plausible account of how dopaminergic genes could control directed and random exploration. More broadly, it has been recognized that dopamine plays a fundamental and multifaceted role in uncertainty computations (Babayan et al., 2018; Costa et al., 2015; Daw et al., 2006a; Friston et al., 2012; Gershman, 2017; Starkweather et al., 2017), raising the important question of whether there exists a deeper principle that ties these computations together.

## Acknowledgments

We are grateful to Athina Tzovara, Eric Schulz and John Mikhael for helpful discussions. This research was supported by the Toyota Corporation and the Office of Naval Research (award N000141712984).

## Footnotes

1 In some applications of the Kalman filter (e.g., Daw et al., 2006b; Speekenbrink and Konstantinidis, 2015), the latent variable is assumed to be time-varying according to a linear-Gaussian dynamical system. However, in our case the reward probabilities are stationary, so we omit dynamics from the learning equations.

2 Note that these criteria take into account model complexity at both the subject-level and group-level.

## References

Auer, P., Cesa-Bianchi, N., Fischer, P. (2002). Finite-time analysis of the multiarmed bandit problem. Machine Learning, 47:235‒256.

Babayan, B. M., Uchida, N., Gershman, S. J. (2018). Belief state representation in the dopamine system. Nature Communications, 9:1891.

Bilder, R. M., Volavka, J., Lachman, H. M., Grace, A. A. (2004). The catechol-o-methyltransferase polymorphism: relations to the tonic-phasic dopamine hypothesis and neuropsychiatric phenotypes. Neuropsychopharmacology, 29:1943.

Blanco, N. J., Love, B. C., Cooper, J. A., McGeary, J. E., Knopik, V. S., Maddox, W. T. (2015). A frontal dopamine system for reflective exploratory behavior. Neurobiology of Learning and Memory, 123:84‒91.

Colzato, L. S., Waszak, F., Nieuwenhuis, S., Posthuma, D., Hommel, B. (2010). The flexible mind is associated with the catechol-o-methyltransferase (comt) val158met polymorphism: evidence for a role of dopamine in the control of task-switching. Neuropsychologia, 48:2764‒2768.

Costa, V. D., Tran, V. L., Turchi, J., Averbeck, B. B. (2015). Reversal learning and dopamine: a bayesian perspective. Journal of Neuroscience, 35:2407‒2416.

Daw, N. D., Courville, A. C., Touretzky, D. S. (2006a). Representation and timing in theories of the dopamine system. Neural Computation, 18:1637‒1677.

Daw, N. D., O’doherty, J. P., Dayan, P., Seymour, B., Dolan, R. J. (2006b). Cortical substrates for exploratory decisions in humans. Nature, 441:876‒879.

Durstewitz, D. Seamans, J. K. (2008). The dual-state theory of prefrontal cortex dopamine function with relevance to catechol-o-methyltransferase genotypes and schizophrenia. Biological Psychiatry, 64:739‒749.

Fienberg, A., Hiroi, N., Mermelstein, P., Song, W.-J., Snyder, G., Nishi, A., Cheramy, A., O’callaghan, J., Miller, D., Cole, D., et al. (1998). Darpp-32: regulator of the efficacy of dopaminergic neurotransmission. Science, 281:838‒842.

Frank, M. J., Doll, B. B., Oas-Terpstra, J., Moreno, F. (2009). Prefrontal and striatal dopaminergic genes predict individual differences in exploration and exploitation. Nature Neuroscience, 12:1062‒1068.

Friston, K., Rigoli, F., Ognibene, D., Mathys, C., Fitzgerald, T., Pezzulo, G. (2015). Active inference and epistemic value. Cognitive Neuroscience, 6:187‒214.

Friston, K. J., Shiner, T., FitzGerald, T., Galea, J. M., Adams, R., Brown, H., Dolan, R. J., Moran, R., Stephan, K. E., Bestmann, S. (2012). Dopamine, affordance and active inference. PLoS Computational Biology, 8:e1002327.

Gershman, S. J. (2017). Dopamine, inference, and uncertainty. Neural Computation, 29:3311‒3326.

Gershman, S. J. (2018a). Deconstructing the human algorithms for exploration. Cognition, 173:34‒42.

Gershman, S. J. (2018b). Uncertainty and exploration. bioRxiv, page 265504.

Ghavamzadeh, M., Mannor, S., Pineau, J., Tamar, A., et al. (2015). Bayesian reinforcement learning: A survey. Foundations and Trends in Machine Learning, 8(5-6):359‒483.

Glimcher, P. W. (2011). Understanding dopamine and reinforcement learning: the dopamine reward prediction error hypothesis. Proceedings of the National Academy of Sciences, 108(Supplement 3):15647‒15654.

Greshake, B., Bayer, P. E., Rausch, H., Reda, J. (2014). opensnp-a crowdsourced web resource for personal genomics. PLoS One, 9:e89204.

Hewitt, J. K. (2012). Editorial policy on candidate gene association and candidate gene-by-environment interaction studies of complex traits. Behavior Genetics, 42:1‒2.

Humphries, M. D., Khamassi, M., Gurney, K. (2012). Dopaminergic control of the exploration-exploitation trade-off via the basal ganglia. Frontiers in Neuroscience, 6:9.

Kakade, S. Dayan, P. (2002). Dopamine: generalization and bonuses. Neural Networks, 15:549‒559.

Krueger, P. M., Wilson, R. C., Cohen, J. D. (2017). Strategies for exploration in the domain of losses. Judgment and Decision Making, 12:104‒117.

Naude, J., Didienne, S., Takillah, S., Prevost-Solie, C., Maskos, U., Faure, P. (2018). Acetylcholine-dependent phasic dopamine activity signals exploratory locomotion and choices. bioRxiv, page 242438.

Niv, Y., Edlund, J. A., Dayan, P., and O’Doherty, J. P. (2012). Neural prediction errors reveal a risk-sensitive reinforcement-learning process in the human brain. Journal of Neuroscience, 32:551‒562.

Onge, J. R. S., Abhari, H., Floresco, S. B. (2011). Dissociable contributions by prefrontal d1 and d2 receptors to risk-based decision making. Journal of Neuroscience, 31:8625‒8633.

Onge, J. R. S., Ahn, S., Phillips, A. G., Floresco, S. B. (2012). Dynamic fluctuations in dopamine efflux in the prefrontal cortex and nucleus accumbens during risk-based decisionmaking. Journal of Neuroscience, 32:16880‒16891.

Schiffmann, S., Desdouits, F., Menu, R., Greengard, P., Vincent, J., Vanderhaeghen, J., Girault, J. (1998). Modulation of the voltage-gated sodium current in rat striatal neurons by darpp-32, an inhibitor of protein phosphatase. European Journal of Neuroscience, (4):1312.

Schulz, E., Konstantinidis, E., Speekenbrink, M. (2015). Learning and decisions in contextual multi-armed bandit tasks. In Proceedings of the 37th Annual Conference of the Cognitive Science Society, pages 2122‒2127.

Schulz, E., Wu, C. M., Ruggeri, A., Meder, B. (2018). Searching for rewards like a child means less generalization and more directed exploration. bioRxiv, page 327593.

Slifstein, M., Kolachana, B., Simpson, E., Tabares, P., Cheng, B., Duvall, M., Frankle, W. G., Weinberger, D., Laruelle, M., and Abi-Dargham, A. (2008). Comt genotype predicts cortical-limbic d1 receptor availability measured with [11c] nnc112 and pet. Molecular Psychiatry, 13:821‒827.

Somerville, L. H., Sasse, S. F., Garrad, M. C., Drysdale, A. T., Abi Akar, N., Insel, C., and Wilson, R. C. (2017). Charting the expansion of strategic exploratory behavior during adolescence. Journal of Experimental Psychology: General, 146:155‒164.

Speekenbrink, M. Konstantinidis, E. (2015). Uncertainty and exploration in a restless bandit problem. Topics in Cognitive Science, 7:351‒367.

Srinivas, N., Krause, A., Seeger, M., Kakade, S. M. (2010). Gaussian process optimization in the bandit setting: No regret and experimental design. In Proceedings of the 27th International Conference on Machine Learning, pages 1015‒1022.

Starkweather, C. K., Babayan, B. M., Uchida, N., Gershman, S. J. (2017). Dopamine reward prediction errors reflect hidden-state inference across time. Nature Neuroscience, 20:581.

Stopper, C. M., Khayambashi, S., Floresco, S. B. (2013). Receptor-specific modulation of risk-based decision making by nucleus accumbens dopamine. Neuropsychopharmacology, 38:715.

Thompson, W. R. (1933). On the likelihood that one unknown probability exceeds another in view of the evidence of two samples. Biometrika, 25:285‒294.

Verharen, J. P., de Jong, J. W., Roelofs, T. J., Huffels, C. F., van Zessen, R., Luijendijk, M. C., Hamelink, R., Willuhn, I., den Ouden, H. E., van der Plasse, G., et al. (2018). A neuronal mechanism underlying decision-making deficits during hyperdopaminergic states. Nature Communications, 9:731.

Warren, C. M., Wilson, R. C., van der Wee, N. J., Giltay, E. J., van Noorden, M. S., Cohen, J. D., Nieuwenhuis, S. (2017). The effect of atomoxetine on random and directed exploration in humans. PloS One, 12:e0176034.

Wilson, R. C., Geana, A., White, J. M., Ludvig, E. A., Cohen, J. D. (2014). Humans use directed and random exploration to solve the explore-exploit dilemma. Journal of Experimental Psychology: General, 143:2074‒2081.

Zajkowski, W. K., Kossut, M., Wilson, R. C. (2017). A causal role for right frontopolar cortex in directed, but not random, exploration. eLife, 6:e27430.

